# Isolation, Detection and Characterization of Aerobic Bacteria from Honey Samples of Bangladesh

**DOI:** 10.1101/298695

**Authors:** Sarbojoy Saha

## Abstract

Honey is a sweet substance made from the nectar of flowers and other chemical secretions from the bees’ bodies who collect nectar from the flowers and bring it to their hives to transform it to the thick, golden and sweet liquid that we call honey. The benefits of honey are not just limited to its basic use as a natural sweetener, but also its medicinal properties. The purpose of this study was to identify the bacteria that are present in honey commonly found in Bangladesh, which can tolerate the antimicrobial conditions of honey and survive in it. Fortunately, such bacteria could be detected, isolated and characterized by morphological and biochemical tests. The predominant type of bacteria commonly found in both raw and commercial honeys of Bangladesh are gram positive cocci such as streptococci, staphylococci, micrococci, bacilli and lactobacilli. Few gram negative bacteria were also isolated like Escherichia coli (8%) and Klebsiella pneumonia (8%) and some gram negative/gram variable Micrococcus luteus *(75%)*. Hopefully, such knowledge would benefit people in the future as they will know more about the microorganisms present in honey and about the safety and quality of the honey they are about to buy or consume.

## Introduction

Honey is a natural sweetening agent produced by honeybees mainly of the species *Apis Mellifera* which collect nectar, the sweet substance found in flowers and mix it with its digestive secretions from a special gland found in their bodies and use it to make honey. Honey has been used as a natural sweetener in place of sugar for thousands of years. Honey is known to have numerous health benefits as it is full of antibacterial substances, vitamins and antioxidants. Honey is often used as a cure for sore throats, colds, cough, dysentery, and several other bacterial infections (Gilliam M and Prest DB, 1985).

Honey is an ancient remedy for the treatment of infected wounds, which has rediscovered by the medical profession, particularly where modern therapeutic agents fail. Aristotle

(384-322 BC), when discussing different honeys, referred to pale honey as being “good as a salve for sore eyes and wounds. (Aristotle, 350 BC) Manuka honey has been reported to exhibit antimicrobial activity against pathogenic bacteria such as *Staphylococcus Aureus (S. Aureus)* and *Helicobacter Pylori (H. Pylori)* making this honey a promising functional food for the treatment of wounds and stomach ulcers (Mandal, Manisha Deb and Shyamapada, 2011).

Honey is a sweetener, has lot of health benefits, however, it contains several microorganisms such as bacteria and fungi such as yeast. The microorganisms found in the honey may come from various sources through different means. These may arise from endogenous sources such as flower nectar, pollen from the bees’ bodies and the digestive secretions from the honeybees. They can also come from external sources such as human handling of honey during its purification process. (Root Al, 1993)

The topic of this research is the identification of bacteria in the honey and these sort of studies have been conducted in Europe. In an Italian study (Sinacori M et al., 2014), the composition of the cultivable microbial populations of 38 nectar honey and honeydew honey samples of different geographical and botanical origin were assessed.

In another study in Italy (Olivieri C et al., 2012) plant, fungal and bacterial DNA in honey specimens were tracked. Snowdon and Cliver (1996) showed that different microbial species in honey may reach a concentration of some thousands forming unit (CFU) per gram. Regarding Italian honeys, lower values of both microbial groups were reported (Piana et al., 1991).

Other studies on this topic such as a study in the United States (Tanzi MG and Gabay MP) focused on the association between honey consumption and infant botulism which is very common there, and another study in Italy (Aureli P et al., 2002) also on honey consumption and infant botulism in Europe. A British study (Grant KA et al., 2013) on Infant botulism advised not to feed honeys to babies and the possible risk factors.

## Composition

Honey primarily contains sugar and water. Sugar accounts for 95–99% of honey dry matter. Majority of these are simple sugars, fructose (38.2%) and glucose (31.3%), which represents 85–95% of total sugars. These are “simple” sugars, 6-carbon sugars that are readily absorbed by the body. Other sugars include disaccharides such as maltose, sucrose, and isomaltose. Few oligosaccharides are also present (White JW, 1957).

Water is the second most important component of honey. Its content is critical, since it affects the storage of honey. The final water content depends on numerous environmental factors during production such as weather and humidity inside the hives, but also on nectar conditions and treatment of honey during extraction and storage (El-leithy MA, El-sibael KB, 1992). Organic acids constitute 0.57% of honey and include gluconic acid which is a by-product of enzymatic digestion of glucose. The organic acids are responsible for the acidity of honey and contribute largely to its characteristic taste (White JW, 1975).

Minerals are present in honey in very small quantities (0.17%) with potassium as the most abundant. Others are calcium, copper, iron, manganese, and phosphorus. Nitrogenous compounds among which the enzymes originate from salivary secretion of the worker honeybees are also present. They have important role in the formation of honey. The main enzymes in honey are invertase (saccharase), diastase (amylase) and glucose oxidase. Vitamins C, B (thiamine) and B2 complex like riboflavin, nicotinic acid and B6 panthothenic acid are also found (Soffer A, 1976).

## Honey production and collection

Knowledge of the process of honey production is important to understand the various ways in which bacteria and other microorganisms may get into the honey. Honey production begins with bees collecting nectar and pollen from flowers but only nectar is used to make honey. Nectar is mostly water with dissolved sugars and the amount of sugars varies greatly but is usually 25–70%. Nectar has been described as a “reward” given by the plant to attract bees (White JW, 1957). Nectar is sucked by honeybees by inserting its proboscis into the flowers nectary and passes it through the oesophagus to the thorax and finally to the abdomen. Pollen is transported back to the hive in the posture pollen baskets on the hind leg whereas the nectar is transported in the stomach. Back in the hive the nectar is placed into wax honeycomb cells and the excess water evaporates until the honey is approximately 83% sugar and 17% water. This takes few days. The cells are then covered with a layer of wax, which is later removed when the bees need to eat the honey. When large amount of nectar is being collected, the bees speed up evaporation by using their wings to ventilate the hive. The sugar is also changed. Sugar in nectar is mostly sucrose, which has large molecules. The bees produce an enzyme (invertase), which breaks each sucrose molecule into glucose and fructose by evaporating the excess water, and converting the sucrose into smaller sugars. The bees therefore make the honey too concentrated for yeast and other microorganisms to grow (Root AL, 1993).

To extract honey, the cap is removed by the use of sharp knife, which has been warmed in hot water. The combs are then put into centrifuge and the honey is removed. Honey is sometimes removed by pressure that necessarily destroys the combs and that may reduce the amount of wax in honey (Bassey IH, 1986).

## Microbes in honey

Microorganisms that survive in honey are those that withstand the concentrated sugar, acidity and other antimicrobial characters of honey. The primary sources of microbial contamination are likely to include pollen, the digestive tracts of honeybees, dirt, dust, air and flowers ‘(Sackelt WG, 1999).

It (Sackelt WG, 1999) has been observed that *Bacillus, Micrococcus and Saccharomyces* species could be readily isolated from honeycombs and adult bees. A number of microbial species have been isolated from the faeces of bee larvae.

*Bacillus* spps are most prevalent followed by Gram -variable pleomorphic bacteria. Mould, Actinomycetes, Gram-negative rods (probably of *Enterobacteriaceae)* and yeast have also been isolated while *Streptomyces* spp were recovered from one larva. This microbial load compares favourably with the intestinal microflora of adult honeybees, which is dominated by Gram variable pleomorphic bacteria of uncertain taxonomic status, *Bacillus* spp,

*Enterobacteriaceae, Penicillium* spp., *Aspergillus* spp., and sometimes frequently *Torulopsis* spp. (Gilliam M, Prest DB.1987). Pollen may be the original source of microbes in the intestines of honeybees (Tysset C et al. 1991). It has been suggested that flowers and hives are more important sources of microbes than the soil. Aerobic spore forming Bacillus were the most frequently encountered microbes on the external surface, crop and intestine of the honey bees (White PB, 1996).

The intestine of bees has been found to contain 1% yeast, 27% Gram-positive bacteria including *Bacillus, Bacteridium, Streptococcus* and *Clostridium* spp; 70% Gram negative or Gram variable bacteria including *Achromobacter, Citrobacter, Enterobacter, Erwinia*, Escherichia coli, *Flavobacterium, Klebsiella, Proteus*, and *Pseudomonas*. The primary sources of sugar tolerant yeast are flowers and soil (Snowdon JA, Cliver DO, 1996).

Secondary sources of microbial contamination in honey are human, equipment, containers, wind, dust etc. Yeasts have been recovered from equipment in honey houses; contaminated equipment can introduce yeast into clean honey (White PB, 1996). Possible routes of transmission into extracted honey would include air (in the house or while the honey was packed), food handlers (from skin infections, sneezing or faecal contamination).

Microorganisms found in honey have been identified. They include bacteria, yeasts and moulds. Most bacteria and other microbes cannot grow or reproduce in honey i.e. they are dormant and this is due to antibacterial activity of honey. Various bacteria have been inoculated into aseptically collected honey held at 20°C. The result showed loss of bacterial viability within 8-24 days. It is only the spore forming microorganisms that can survive in honey at low temperature. The spore count remained the same 4 months after. *Bacillus cereus, Clostridium perfringes* and *Clostridium botulinum* spores were inoculated into honey and stored at 25°C. The *Clostridium botulinum* population did not change over a year at 4°C. At 65°C however, no spores were found after 5 days of storage. It has been observed that if honey is diluted with water, it supports the growth of non-pathogenic bacterial strains and killing of dangerous strains. Solution of less than 50% honey in water sustained bacterial life for longer periods but never exceeding forty days.

## Objective/Aim of the project

The aim of the project is to isolate the bacteria from honey samples and identify the bacteria by morphological and other biochemical techniques. A total of thirteen honey samples were tested, six commercial and the other seven raw honeys. The bacteria isolated were identified and characterized by the methods described in the methodology.

## Materials and methods

### Flow chart of the research steps and activities

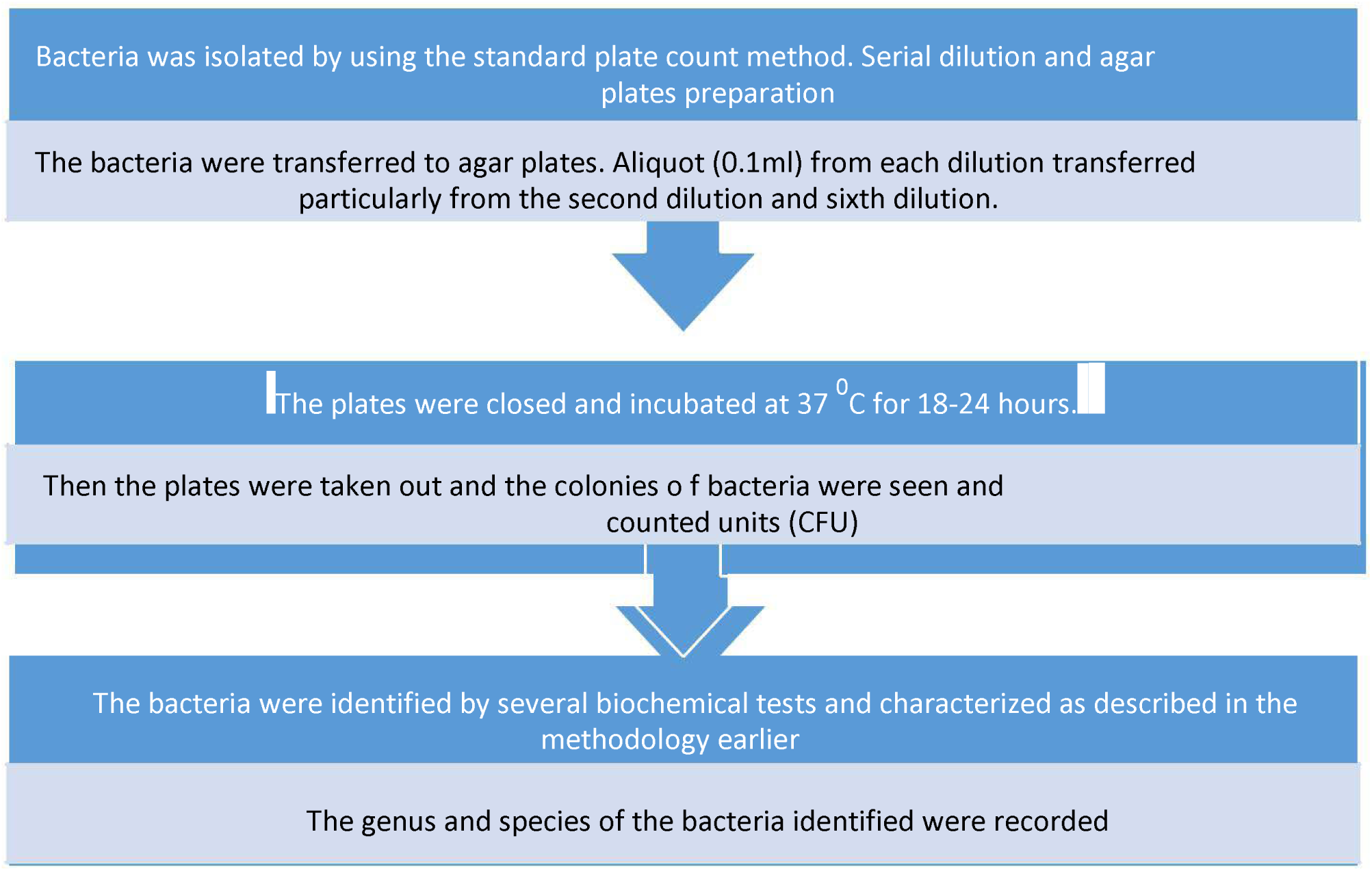

### Isolation of bacteria

Bacteria was isolated using the standard plate count method. Serial dilution and gram staining was done. The aliquots from each dilution were transferred to the plates and spread plated. Then, the plates were incubated for a day. The following day sufficient growth was seen in the plates and colony forming units were counted. The bacteria were identified using biochemical tests and staining.

### Serial dilution

For serial dilution, 9 ml of diluent or distilled water was pipetted into six test tubes for making three-fold dilutions. Then 1ml of sample have to be taken by a micropipette and added to the Tube 1 to make one-fold dilution or 10^−1^ dilution. Again 1ml was taken from Tube 1 and transferred to Tube 2 to make second-fold dilution or 10^−2^ dilution, and 1ml was taken from Tube 2 and transferred to Tube 3 to make three-fold dilution or 10^−3^ dilution and so forth. Thus, three-fold dilutions were prepared in this way.

An aliquot of 0.1 ml from each dilution will be transferred to the prepared agar plates, particularly from the second and third dilutions. For each dilution two or three duplicates were done. The plates were incubated at 37 C for 18-24 hours normally. After incubation the plates were taken out for observation of the bacterial colonies which should be formed.

The colonies were observed and the colony forming units(CFU) were counted (Ibrahim AS, 1980).

### Gram staining

For gram staining, at first, a drop of water was placed on a glass slide by a pipette. Inoculating loop was sterilized by a Bunsen flame, and the work surface sterilized by alcohol.

- Then using the sterile loop after it had been flamed until orange, a single bacterial colony was transferred from the agar plate to the drop of water on the slide and spread to make a smear.
- It then was dried for 10 minutes, and heat fixed by a Bunsen flame.
- First stain added was Crystal violet and allowed to stain for 1 minute. After staining for 1 min, the slide was washed with sterile water to remove the stain.
- Then, Gram’s Iodine was added by a few drops and allowed to stain for 1 minute. After staining, slide was rinsed by water to remove as much iodine as possible.
- Then, ethanol or decolorizer was added and the slide washed with water later.
- Safranine addition at the end as a counter stain, and slide was washed. Then slide was observed under a microscope. Gram+ bacteria give purple colour, Gram – pink.

### Biochemical tests

Biochemical tests were carried out for identification of the bacteria. Several tests such as catalase test, oxidase test, coagulase, mannitol salt agar plating, Simmon’s citrate agar, EMB agar, motility agar, methyl red-voges proskauer broth were carried out, the results of which are shown below.

**Table 1:** Biochemical assay based characterization of bacteria from honey samples. (Naseer, S et al.2015)

| Bacterial isolate s | Staphylococcus aureus | Micrococcus luteus | Streptococcus pyogenes | Streptococcus faecalis | Streptococcus epidermidis | Corynebacterium sp. | Klebsiella pneumonia | Escherichia coli | Salmonella sp | Proteus sp |
| --- | --- | --- | --- | --- | --- | --- | --- | --- | --- | --- |
| Catalase | + | + | - | - | - | + | + | + | - | + |
| Oxidase | - | + | - | - | - | - | - | - | - | - |
| Coagulase | + | - | - | - | - |  |  |  |  |  |
| Motility | - | - | - | - | - | - | - | + | + | + |
| Indole | - | - | - | - | - | - | - | + | - | + |
| Ethyl red | + | - | - | - | - | - | - | + | + | + |
| VP test | + | - | - | - | - | - | - | + | + | + |
| Citrate test | + | - | - | + | - | - | + | - | + | - |
| Growth on MacConkey | - | - | - | + | - | + | + | + | + | + |
| Hemolysis on blood agar | B | y | B | y | y | y | y | a | y | A |
| Oxygen requirement | F.anaerobe | O.aerobe | F.anaerobe | F.anaerobe | Aerobe | F.anaerobe | F.anaerobe |  | F.anaerobe | F.anaerobe |
| Lactose fermentation | + | - | - | + | + | - | + | + | - | - |
| Sucrose fermentation | + | - | - | + | + | - | + | + | - | - |

## Results

**Table 2:** The bacteria isolated and detected from the seven raw honey samples.

| Location and flower type | Bacteria identified | Colony forming units |  |  | Mean count |
| --- | --- | --- | --- | --- | --- |
|  |  | 10-1 | 10-2 | 10-3 |  |
| Jessore mixed honey | Pathogenic: *Staphylococcus aureus* and *Streptococci*<br>Non-pathogenic: *Micrococcus luteus* | 3 | 3 | 3 | 3 |
| Rajshahee lychee honey | Pathogenic: *Bacillus cereus/bacilli*<br>Non-pathogenic: *Micrococcus luteus* | >800 | 400 | 88 | >200 |
| Rajshahi mustard honey | Pathogenic: *Staphylococcus aureus, streptococci* and *bacilli*<br>Non-pathogenic: *Micrococci* | 2 | 2 | 0 | 2 |
| Khulna khalsi flower honey | Pathogenic: *Staphylococci* and *streptococci*<br>Non-pathogenic: *Lactobacilli, micrococci* | 8 | 5 | 5 | 5 |
| Khulna goran flower honey | Pathogenic: *Staphylococcus aureus* and *streptococci*<br>Non-pathogenic: *Lactobacilli, micrococci* | 18 | 18 | 18 | 18 |
| Chapai' Nawabganj kul flower honey | Pathogenic: *Streptococci*<br>Non-pathogenic: *Micrococci* | 9 | 7 | 5 | 7 |
| Chapai' Nawabganj black cumin flower honey | Pathogenic: *Streptococci, E. coli*<br>Non-pathogenic: *Micrococci* | 4 | 4 | 2 | 4 |

**Table 3:** The bacteria isolated from the six commercial honey samples, along with their colony count.

| Name of honey sample | Bacteria identified | Colony forming units |  |  | Mean colony count |
| --- | --- | --- | --- | --- | --- |
|  |  | 10-1 | 10-2 | 10-3 |  |
| Dabur honey | *Pathogenic: Streptococci, staphylococcus aureus and bacilli.*<br>*Non-pathogenic: Micrococci* | 5 | 5 | 5 | 5 |
| AP Black cumin honey | *Pathogenic: staphylococcus aureus, streptococci*<br>*Non-pathogenic: Micrococci* | 100 | 48 | 8 | 5 |
| AP mixed honey | *Pathogenic: Klebsiella, streptococci, staphylococcus aureus*<br>*Non-pathogenic: Micrococci* | 4 | 4 | 1 | 4 |
| Musaffa multi-floral honey | *Pathogenic: Streptococci, staphylococcus aureus*<br>*Non-pathogenic: Micrococci* |  |  |  | 78 |
| Musaffa khalsi flower honey | *Pathogenic: Streptococci, staphylococcus aureus*<br>*Non-pathogenic: Micrococci* | 17 | 16 | 16 | 16 |
| Proshika mustard honey | *Pathogenic: Streptococci, staphylococcus aureus*<br>*Non-pathogenic: Micrococci* | >200 | 6 | 6 | 70 |

**Table 4:** The similarities and differences between this project and studies conducted elsewhere.

| Name of other study | Similarities | Differences |
| --- | --- | --- |
| Assessment of Microbial Load of un-pasteurized fruit juices and in vitro antibacterial potential of honey against bacterial isolates (Iqbal, MN et al.2015) | Both the studies involved the bacteria *Bacilli*, *Staphylococcus aureus*, *Klebsiella* and *E.coli* | But the other study involved the bacteria *Pseudomonas* and *Enterobacter*. This study did not. |
| Antibacterial efficacy of raw and commercially available honey<br>(Rahman S, Salehin F, Iqbal A,2011) | Both the studies involved *Staphylococcus aureus*, and *E.coli*. | But the other study also used the bacteria *Shigella* spp.,*Pseudomonas aeruginosa*, *Salmonella* while this study did not. |
| Honey: a reservoir for microorganisms and an inhibitory agent for microbes<br>Peter B. Olaitan et al.,2007) | Both studies mentions bacilli, *Micrococcus*, *Streptococcus*, *E.coli* and *Klebsiella* among others. | But the other study also detected yeast, *Streptomyces*, *Mould*, *Actinomycetes* and *Enterobacteriaceae* and others unlike this study done in Bangladesh. |
| Identification and roles of non-pathogenic microflora (Gilliam M,1997) | Similarities with this project include the detection of bacillus species and *Enterobacteriaceae*. | Differences were yeast and gram variable pleomorphic bacteria not detected in this local study in Bangladesh. |
| Microorganisms in honey.(Snowdon JA, Cliver DO,1996) | Both the studies identified *bacilli*. | But the study done by Snowdon and Cliver also detected yeast, *coliform* and spore forming *clostridium* |

Similar tests were done for the other six commercial honey samples mentioned later. The results obtained have been already summarized earlier in the table, both the species identified and colony counts.

1. In the Jessore mixed honey, the bacteria isolated were *Micrococcus luteus, staphylococcus aureus* and *Streptococus*. And the colony count was two-three CFUs on average on each dilution plate.
2. In the Rajshahi lychee honey, *Micrococcus luteus* and *Bacillus cereus* were isolated and identified. The first dilution plate had such numerous growth that it was uncountable. The second had a little less but still a lot of colonies at four hundred CFUs. The third plate had the least at eighty-eight CFUs.
3. The mustard honey from Rajshahi had *Micrococcus luteus, Staphylococcus aureus and Bacillus cereus*. After serial dilution and spread plating, two CFUs were seen on the first and second dilution plates and none on the third.
4. Using the Khulna Khalsi flower honey, the bacteria identified were *Micrococcus luteus, Staphylococus aureus, Streptococcus and Lactobacillus acidophilus*. The spread plates made earlier gave eight colonies in the first dilution plate and five colonies each in the second and third dilution plate.
5. The Khulna Goran flower honey also had the same species of bacteria like the khalsi flower honey from Khulna. Eighteen CFUs were observed on each dilution plate on average after incubation for a day.
6. The Kul flower honey from Chapai’Nawabganj had *Micrococcus luteus and Streptcoccus*. The first dilution plate had nine colonies and the second had seven CFUs. The third dilution plate had five.
7. The Chapai’Nawabganj Black cumin honey had *Micrococcus luteus, Streptococcus and Escherichia*

**Fig 1:**
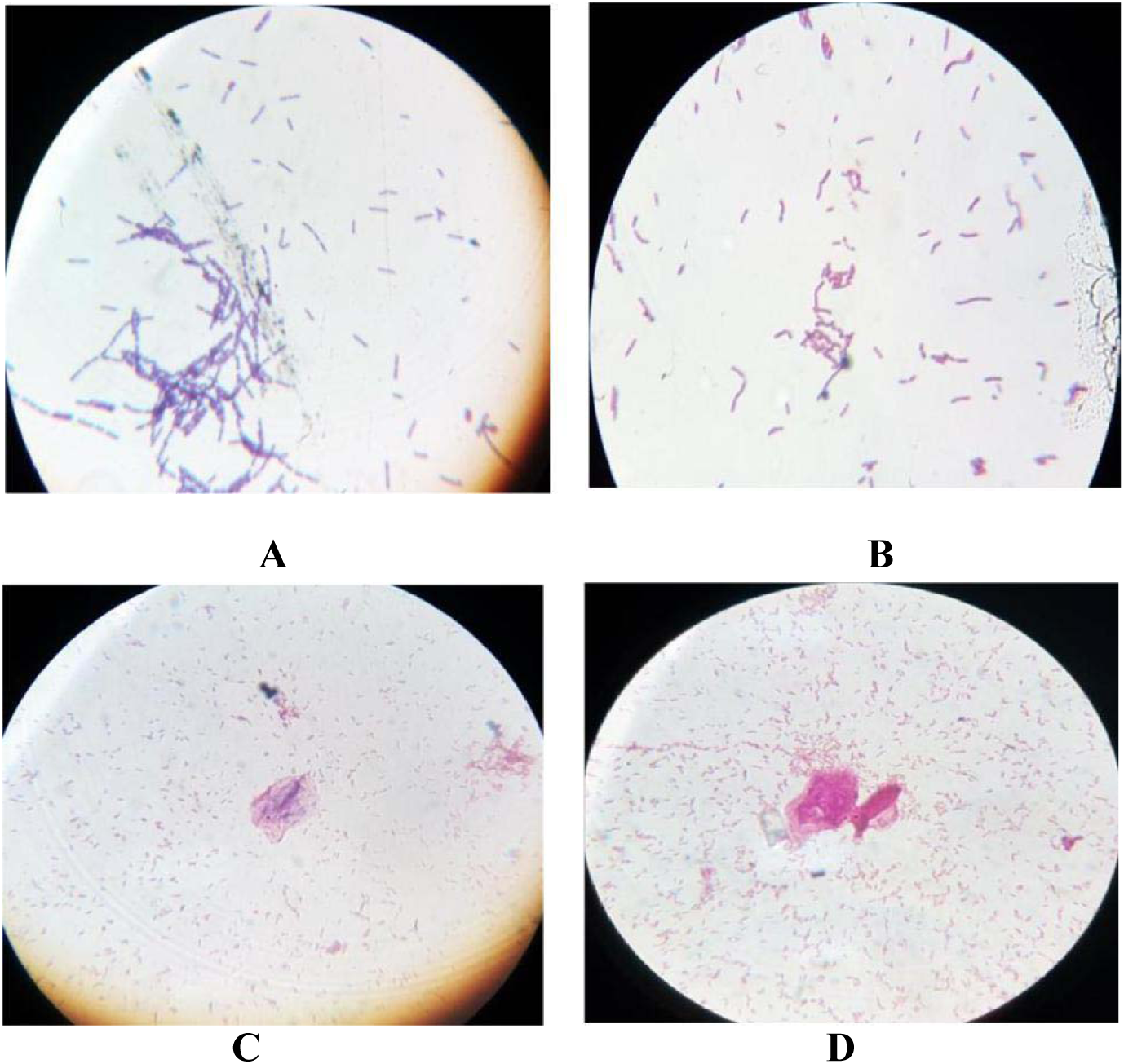
gram staining photos showing gram-positive *Bacilli* found in Rajshahi lychee honey (A), gram-positive *Lactobacilli* found in Khulna goran flower honey (B), gram-negative *E. coli* found in Chapai’Nawabganj black cumin flower honey (C), gram-negative *Klebsiella* found in AP mixed honey (D).

**Fig 2:**
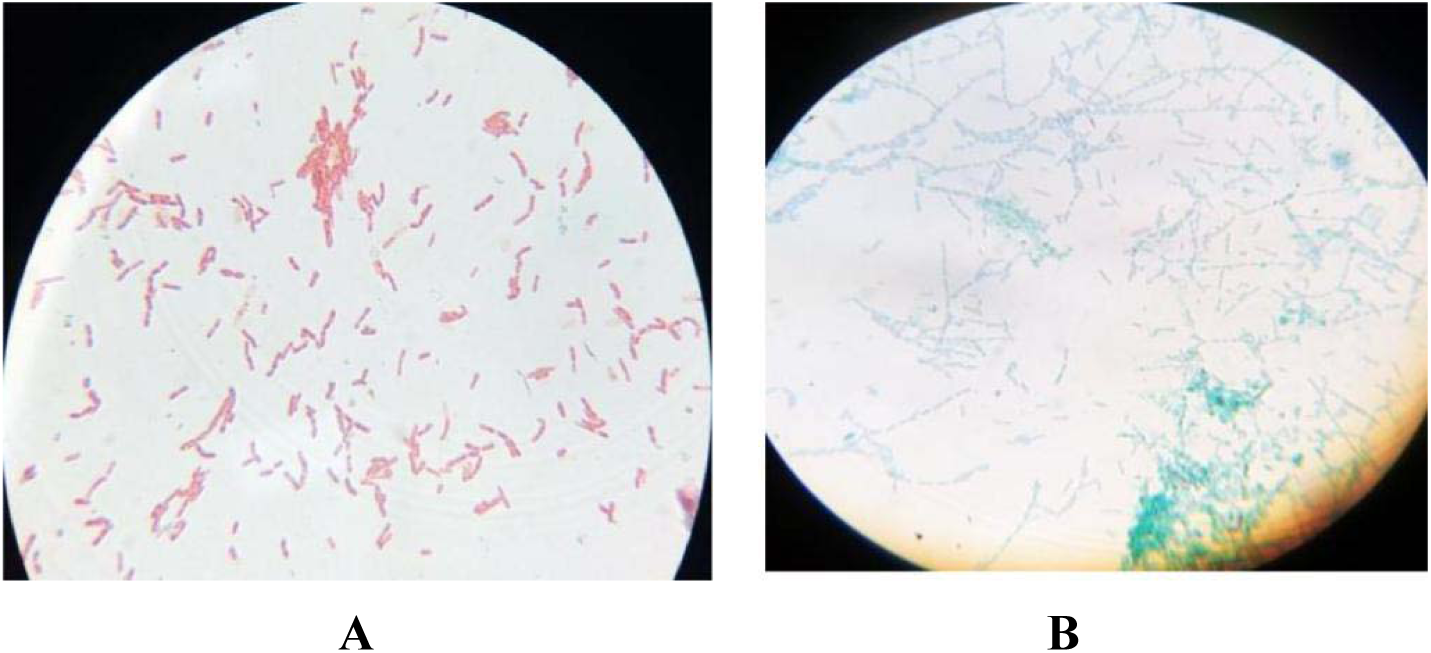
The spore staining photos showing non-spore forming *lactobacilli* in red found in Khulna goran flower honey (A) and endospore forming *bacilli* in green found in Rajshahi lychee honey (B)

### Results from the six commercial honey samples

There were a total of six commercial honey samples from different brands which were collected and tested.

The bacteria isolated, identified and characterized from the commercial honey samples are also listed below with their colony count.

1. Using the Dabur honey sample *Micrococcus luteus, Staphylococcus aureus, Streptococcus and bacillus cereus* were identified. Five colonies/CFUs were seen on the first and second dilution plates.
2. Using the commercial AP Black cumin honey the bacteria identified were *Micrococcus luteus, Staphylococcus aureus and Streptococcus*. A hundred colonies/CFUs were observed on the first dilution plate and forty-eight CFUs on the second dilution plate. The third had seven CFUs.
3. The commercial AP mixed honey had *Micrococcus luteus, Staphylococcus aureus, Streptococcus and Klebsiella pneumonia*. The following day, the plates were taken out and four CFUs were observed on the first and second dilution plate and one CFU on the third.
4. The bacteria identified after isolation from the Musaffa khalsi flower honey were *Micrococcus luteus, Staphylococcus aureus and Streptococcus*. Seventeen CFUs were present on the first dilution plate and sixteen CFUs on the second and third dilution plates.
5. The Musaffa multi-floral honey gave similar species of bacteria as the Musaffa khalsi flower honey. The colony count in the first dilution plate was numerous or uncountable. The second dilution plate had thirty CFUs and the third one had five.
6. Using the Proshika mustard honey, the bacteria identified were *Micrococcus luteus, Staphylococcus aureus and Streptococcus*. Numerous growth was seen on the first dilution plate, so much that it was uncountable (> two hundred CFUs). The second and third dilution plates had six CFUs each.

## Discussion

The aim or objective of this research project was to isolate and identify the bacteria which are present in the honey of Bangladesh. Both raw and commercial honey samples were used for this purpose. There were a total of thirteen honey samples that were tested. Six of them were commercial honey samples, and the other seven were raw honey samples collected from different parts of the country. Then, these were brought and stored at home in a cool, dry place before laboratory use.

The bacteria isolated from the honey samples were grown and maintained on nutrient agar, and were identified by morphological and biochemical techniques. At first, only the raw honey samples were tested and the bacteria present in those honey samples were isolated and identified. This was followed by the testing of the commercial honey samples. Mostly, the bacteria isolated and identified were *micrococci, streptococci and staphylococci* as are commonly found in honey. But, there were also a few exceptions like *lactobacilli* found in the Khulna khalsi flower and goran flower honey. And *E. coli* found in the Chapai’Nawabganj Black cumin flower honey. A similar study was done with Argentinean honeys (Iurlina MO, Fritz R.2005).

Several studies of this kind have been carried out in Europe like in Italy and England. In Italy, a similar study was carried out (Olivieri C et al., 2012) where plant, fungal and bacterial DNA in honey specimens were tracked. Snowdon and Cliver (1996) showed that different microbial species in honey may reach a concentration of some thousands forming unit (CFU) per gram. Whereas in Italian honeys, lower values of both microbial groups were reported (Piana et al., 1991).

The drawback of this process was that the raw honey samples were especially difficult to collect, as they were from parts of the country and also they were quite expensive. Another limitation was a shortage of petri dishes in the laboratory at first, and a shortage of test tubes. A sudden deficiency of nutrient agar also occurred in the laboratory adding to the drawbacks.

The project gives us information in detail about the types of bacteria found in the honey of Bangladesh. Hopefully, this information will enable the general public to be more aware of what type of honey they are buying from the markets and consuming, and they will be able to make more informed choices after knowing this information found and presented from this research project. And common digestive problems which are often experienced after consumption of honey and products made with honey can also be avoided in the future if the people have a greater awareness about this topic. And, such a research project will encourage students and researchers to be more curious about honey and the microorganisms which commonly live in it, and more research projects of this kind will be undertaken in the near future. Though honey has antibacterial properties, still some bacteria can survive and grow in it. In the future, the mechanism of action of some bacteria which

